# Diverse Phosphoserine Phosphatases Exhibit Maximum Activity at an Intermediate Binding Affinity in Accord with the Sabatier Principle of Catalysis

**DOI:** 10.1101/2023.03.10.532031

**Authors:** Yoko Chiba, Hideshi Ooka, Marie Wintzer, Nao Tsunematsu, Takehiro Suzuki, Naoshi Dohmae, Ryuhei Nakamura

**Author notes:** **Corresponding Authors:** Yoko Chiba, Hideshi Ooka, **Email:** /.

## Abstract

A unified framework to rationalize enzymatic activity is essential to understand cellular function and metabolic evolution. Recent studies have shown that the activity of several hydrolases is maximized when the substrate binding affinity (Michaelis-Menten constant: *K_m_*) is neither too strong nor too weak. This is because an intermediate *K_m_* resolves the trade-off between *K_m_* and *k_cat_*, in accord with the Sabatier principle of artificial catalysis. However, it remains unclear whether this concept is applicable to enzymes in general, especially for those which catalyze the same reaction but have evolved under different selection pressures due to the phylogeny or physiology of the host organism. Here, we demonstrate that the activity of 10 distinct wild-type phosphoserine phosphatases (PSP) exhibits a maximum at an intermediate binding affinity (*K_m_* ≈ 0.5 mM), indicating that they also follow the Sabatier principle. Furthermore, by considering not only *K_m_* but also the equilibrium rate constant 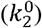 of each enzyme, we have succeeded in rationalizing the PSP activity quantitatively. 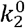 is the rate constant of product release (ES → E + P) in the absence of any driving force, and a large 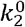 allows *k_cat_* to be increased without increasing *K_m_*. Although the traditional Sabatier principle considers only the binding affinity (*K_m_*), we show that the additional contribution of 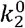 drastically improves the consistency between experiments and theory. Our expanded framework which quantitatively explains the activity of phylogenetically and physiologically diverse enzymes with respect to their physicochemical parameters may lead to the rational design of highly active enzymes.

## INTRODUCTION

Enzymes are the primary component of biological metabolism and understanding how their activity is determined is of prime importance, not only for understanding the design principles underlying metabolic networks in nature, but also to promote biotechnological applications including rational design of enzymes and metabolisms. Historically, their reaction rate (*v*) has been rationalized by the Michaelis-Menten equation (Eq.1) (1–4) based on the reaction mechanism shown in Eq. 2:

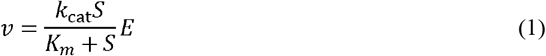

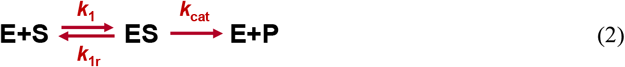

Here, the reaction rate is expressed as a function of a rate constant *k_cat_*, the Michaelis-Menten constant (*K_m_*), and the substrate (S) and enzyme (*E*) concentrations. *K_m_* can be interpreted as the substrate binding affinity of the enzyme and is defined as:

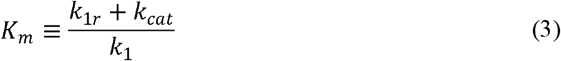

*K_m_* and *k_cat_* are obtained routinely by evaluating the reaction rate at various substrate concentrations and then fitting the experimental data with the Michaelis-Menten equation. Historically, enzymes with small *K_m_* and large *k_cat_* were considered to be good catalysts due to their high reaction rate (Eq. 1), and therefore, 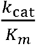 has been used to evaluate catalytic efficiency (5). However, there are still no widely accepted guidelines on the values of *K_m_* and *k_cat_* which can maximize the reaction rate.

Recently, several studies have provided critical insight on the optimum *K_m_* value (6, 7). Kari *et al* have shown experimentally that the activity of cellulases and poly (ethylene terephthalate) (PET) hydrolases can be maximized when the *K_m_* is neither too large nor too small (6, 8, 9). In addition, Dyla *et al* have also reported that kinase activity is maximized at an intermediate docking affinity (*k*_off_) between the enzyme and the substrate (10). These findings are consistent with the Sabatier principle in artificial catalysis (11, 12), which states that there is an optimal binding affinity between a catalyst and its substrate. This principle was originally proposed in heterogeneous catalysis, but its applicability has recently been expanded to homogeneous catalysis (12–14). The gradual expansion of the Sabatier principle from heterogeneous catalysis to homogeneous catalysis and heterogeneous enzymes suggests that tuning the binding affinity may be a general strategy to increase the activity of both enzymes and artificial catalysis. However, the broad applicability of the Sabatier principle to enzymes is still unclear, due to the scope in which the relationship between activity and *K_m_* has been analyzed so far. In the case of PET hydrolase, two enzymes from the same bacterium were used and the *K_m_* was manipulated by adding different concentrations of a surfactant. A larger variety of enzymes (36 wild-type and 47 mutants) were analyzed in the case of cellulase, although all of them were derived from Fungi. Therefore, the existence of an optimum *K_m_* and the applicability of the Sabatier principle to enzymes in general is still unclear, especially for enzymes which catalyze the same reaction but exist in both phylogenetically and physiologically diverse organisms, which may have evolved under different selection pressures.

Here in this study, we demonstrate that phosphoserine phosphatase (PSP, EC:3.1.3.3) obtained from diverse physiological and phylogenetical backgrounds also exhibits maximum activity at an intermediate *K_m_*. Furthermore, we show that the Sabatier principle can quantitatively rationalize the activity with respect to the *K_m_* after considering differences in the equilibrium rate constant 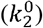 of each enzyme. These results suggest that the expansion of the Sabatier principle presented in this work can provide a unified framework to understand enzymatic activity.

## RESULTS

### Selection of PSPs with various physiologic and phylogenetic background

To assess the possibility of a framework to rationalize enzyme kinetics, PSP was selected as a model enzyme due to its widespread distribution in all three domains of life: Archaea, Bacteria, and Eukaryota. Furthermore, there are three types of PSPs with no clear homology between them, allowing the activity to be compared between enzymes with completely different amino acid sequences. Here we named them type 1, 2 and 3 according to their order of discovery. Type 1 and type 3 PSPs belong to halo acid-like hydrolase superfamily while type 2 PSP belongs to histidine phosphatase superfamily. Type 1 PSP is distributed in all three domains of life: Archaea, Bacteria, and Eukaryota (15–19). In contrast, type 2 and type 3 PSPs are only distributed in Bacteria (20–23). Therefore, the three type 1 PSPs in this study were sampled from Archaea, Bacteria, and Eukaryota, while the four type 2 PSPs and three type 3 PSPs were obtained from Bacteria (Table 1, Supporting information I). Even within the same type, amino acid sequence identities were 32%-55%. The host organisms are also diverse in their physiology. Namely, their optimal growth temperatures ranged from room temperature to 85□, and both autotrophs and heterotrophs were included. Therefore, the diverse set of enzymes chosen in this study provides a stringent criterion to validate the applicability of a framework to rationalize enzymatic activity.

**Table 1.** Origin of PSPs used for this project and the kinetic parameters at 40ĨD.

| Host organism | Domain | Optimal growth Temp (°C) | Protein ID | PSP type | $K_m$ (mM) | $k_{cat}$ (s<sup>-1</sup>) | $k_{cat}/K_m$ |
| --- | --- | --- | --- | --- | --- | --- | --- |
| *Methanocaldococcus jannaschii* | Archaea | 85 | MJ_1594 | 1 | 0.36 | 5.2 | 14.4 |
| *Homo sapiens* | Eukarya | - | PSPH | 1 | 0.008 | 7.4 | 971.8 |
| *Escherichia coli* | Bacteria | 37 | JW4351 | 1 | 0.47 | 66.8 | 142.2 |
| *Hydrogenobacter thermophilus* | Bacteria | 70 | HTH_0103 | 2 | 9.64 | 3.8 | 0.4 |
| *Persephonella marina* | Bacteria | 55-80 | PERMA_0343 | 2 | 1.98 | 7.5 | 3.8 |
| *Chloroflexus aurantiacus* | Bacteria | 50-60 | Caur_0633 | 2 | 0.18 | 2.4 | 13.1 |
| *Thermosynechococcus elongatus* | Bacteria | 50 | TLR1532 | 2 | 0.10 | 2.9 | 29.6 |
| *Thermus thermophilus* | Bacteria | 70 | TT_C1695 | 3 | 0.47 | 11.2 | 23.7 |
| *Meiothermus ruber* | Bacteria | 60-65 | K649_13560 | 3 | 0.55 | 66.7 | 121.3 |
| *Bacillus subtilis* | Bacteria | 25-35 | BSU28940 | 3 | 1.14 | 71.1 | 62.6 |

Type 1 PSP (also called metal-dependent dPSP) and type 3 PSP belong to the haloacid dehalogenase (HAD) like-hydrolase superfamily while type 2 PSP (also called metal-independent iPSP) belong to the histidine phosphatase superfamily.

### Determination of Michaelis-Menten constants

The PSPs were expressed in *E. coli* and purified using FPLC (Fig. S1). No artificial tags were used, because the terminal sequences of some PSPs are known to be critical for their activity (21). The catalytic activity of each enzyme with respect to the substrate concentration is shown in Fig. 1 and Fig. S2. Each enzyme was confirmed to follow Michaelis-Menten kinetics and *K_m_* and *k_cat_* were estimated using non-linear regression (Table 1). The minimum and maximum *K_m_* within this dataset were 0.008 mM (PSPH) and 9.6 mM (HTH_0103), respectively.

**Fig. 1.**
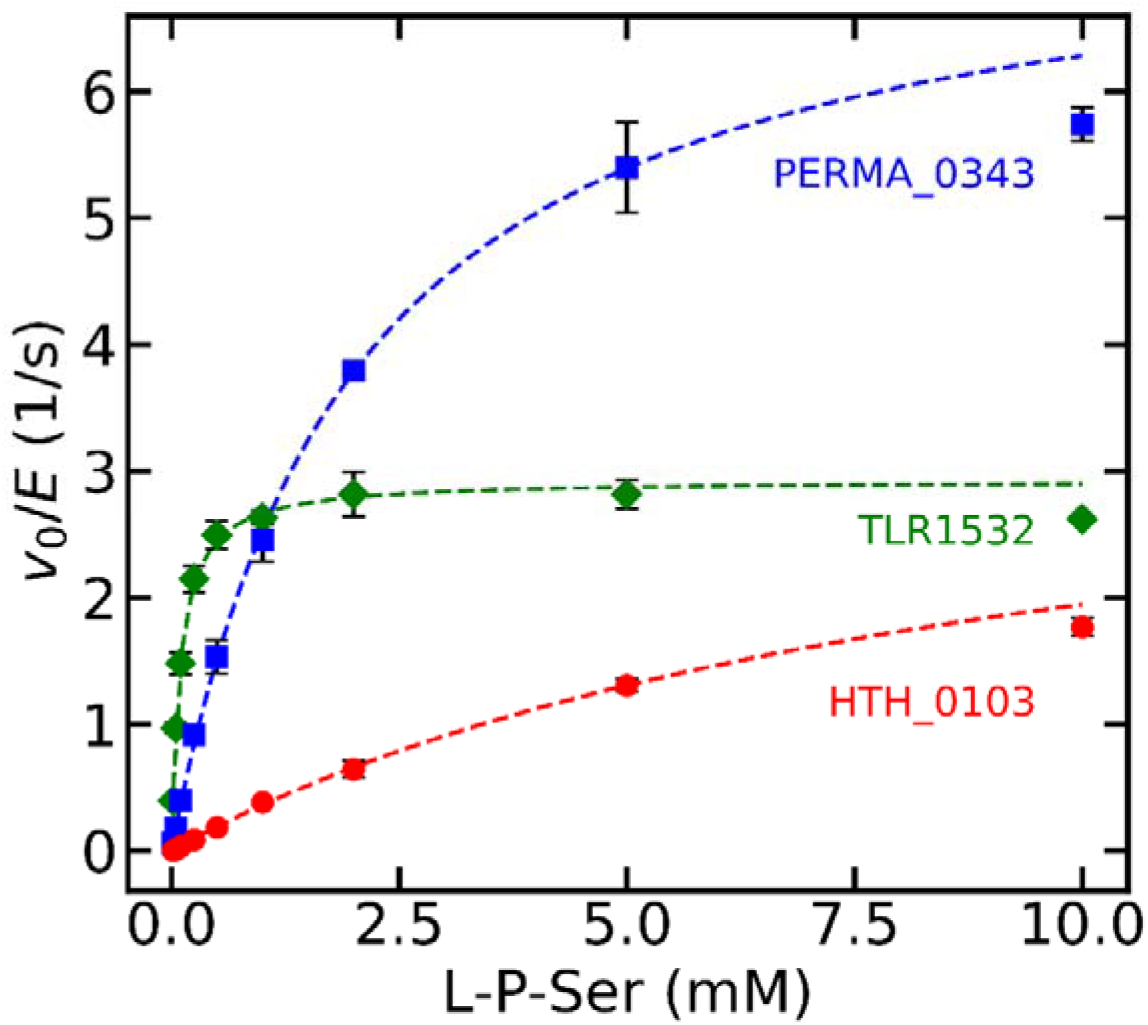
Representative Michaelis-Menten plots of PSPs. The PSP activity from *Thermosynechococcus elongatus* TLR1532 (green diamond), *Persephonella marina* PERMA_0343 (blue square), and *Hydrogenobacter thermophilus* HTH_0103 (red circle) are shown. Error bars indicate their standard deviation.

### Relationship between *K_m_* and PSP activity

The initial reaction rates (*v*_0_/*E*, specific rate (s^-1^)) of PSP showed a local maximum around *K_m_*= 0.5 mM when the reaction rate was plotted against the logarithm of *K_m_* (Fig. 2). The existence of a local maximum is consistent with the case of cellulases and PET hydrolases (6, 8, 9), and is also conceptually consistent with the Sabatier principle which predicts the existence of an optimum binding affinity. However, there are some quantitative discrepancies between the experimental results and the theoretical predictions (24–26) of the Sabatier principle. First, increasing the *K_m_* from 0.008 to 0.1 mM decreases the activity, despite the *K_m_* value approaching the optimum (*K_m_* = 0.5 mM). This is in contrast to the Sabatier principle which predicts the existence of only one local maximum. Furthermore, the *K_m_* which maximized the activity did not shift depending on the substrate concentration, despite previous theoretical models predicting a shift towards larger *K_m_* (24, 25). The physical origin of the shift is because the substrate participates in only the first step of the reaction. When the substrate concentration is increased, the rate of the first step (E + S → ES) would be enhanced, while that of the second step (ES → E + P) would remain constant. Therefore, as long as the enzyme follows Michaelis-Menten kinetics, the optimum *K_m_* should be dependent on the substrate concentration. The results reported for cellulases are in better agreement with theoretical predictions, as there was only one optimal *K_m_* which increased by about two orders of magnitude upon increasing the substrate concentration from 1 to 100 g/L (8). The deviation of PSP from theoretical predictions, such as the presence of multiple local maxima which are insensitive to the driving force, indicates that our more diverse dataset cannot be rationalized quantitatively using the Sabatier principle unless additional factors are considered.

**Fig. 2.**
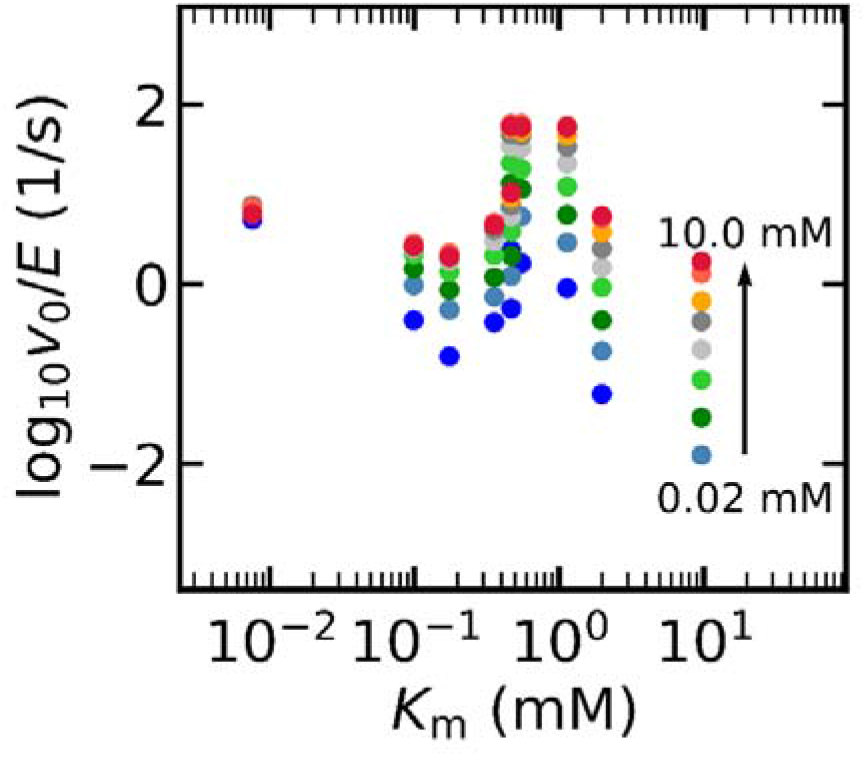
Enzymatic activity (*v*_0_) plotted with respect to the of each enzyme. Each color corresponds to a different substrate concentration (0.02. 0.05, 0.1, 0.25, 0.5, 1, 2, 5 and 10 mM, respectively).

### Theoretical evaluation of enzyme parameters

To assess whether the PSP activity can be discussed within the framework of the Sabatier principle, we have attempted to clarify the origin of the deviation by evaluating the physicochemical parameters of each enzyme. In particular, we have attempted to characterize the substrate binding affinity using the Gibbs free energy (Δ*G*_1_), because it is the standard metric used to apply the Sabatier principle in catalysis. However, although 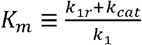 and 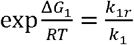 are related, there is no direct method to convert *K_m_* to Δ*G*_1_. Therefore, we used the following equation to evaluate the Δ*G*_1_ from the experimental data of *K_m_* and *k*_2_:

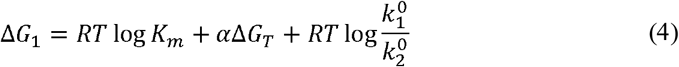

This equation was obtained as described in the Materials and Methods, and allows the binding affinity Δ*G*_1_ to be calculated based on experimentally accessible parameters. A Δ*G_T_* value of −50 kJ/mol was obtained from eQuilibrator after considering concentrations of ions and substrates used in the experiment. *a* is the Bronsted-Evans-Polanyi (BEP) coefficient, and indicates the relationship between activation barriers and the applied driving force. As 0 < *α* < 1, a median value of 0.5 was used, following convention in heterogeneous catalysis. Different values of *α* do not influence the main conclusions as shown in Fig. S4. The ratio 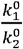 was evaluated based on the Arrhenius prefactors of *k*_1_ and *k_cat_*, which were determined by their temperature dependence (see Material and Methods and the Python code for details). The obtained parameter values are self-consistent in the sense that the *K_m_* value calculated from the estimated parameters are consistent with the experimental value (Fig. S4).

Fig. 3 shows the enzymatic activity with respect to *K_m_* (Fig. 3A,C) and Δ*G*_1_ (Fig. 3B,D). Markers indicate the experimental data, and dashed lines indicate the theoretical activity calculated from the Michaelis-Menten equation (Eq. 3) using kinetic parameters (*K_m_* and *k_cat_*) which satisfy Eqs. 9 and 10. In order to take the relationship between *K_m_* and *k_cat_* into account, we have set the value of 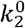 to the average value obtained in the experiments and then calculated the corresponding *k_cat_* at each *K_m_* to draw the theoretical line. Therefore, all parameters were directly calculated based on a physical model, and no parameters were “fitted” to the experiments. In both Fig. 3A and B, there is a clear deviation between the experimental data and the theoretical curve, indicating that the difference between *K_m_* and Δ*G*_1_ is not the main reason why PSP activity deviates from theoretical predictions. On the other hand, the analysis above revealed that each PSP had a markedly different 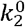 which ranged from 4.32 x 10^-3^ to 3.46 x 10^-1^ s^-1^ (Supporting information) while the theoretical curves were calculated assuming only a single value of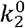. As the two order of magnitude difference in 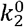 directly corresponds to an activity difference of the same magnitude, we normalized the experimental reaction rates of each enzyme with their 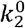 values (Fig. 3C, D). After normalization, the experimental results become consistent with the theoretical curve. Namely, there is only one binding affinity (either *K_m_* or Δ*G*_1_) which maximizes the normalized activity and the optimal binding affinity shifted to higher concentration upon increasing the substrate concentration. This shift is reasonable considering that if *S* » *K_m_*, enzymatic activity is saturated at *V_max_* and decreasing *K_m_* does not increase activity. On the other hand, decreasing *K_m_* may be detrimental because it tends to decrease *k_cat_* based on the definition shown in Eq. 3. Indeed, Kari et al have shown that *K_m_* and *k_cat_* are highly correlated in cellulases (r^2^ = 0.95 in the log scale) (8). Therefore, at larger substrate concentrations, a larger *K_m_* is preferable, leading to a concentration-dependent shift of the optimum *K_m_*.

**Fig. 3.**
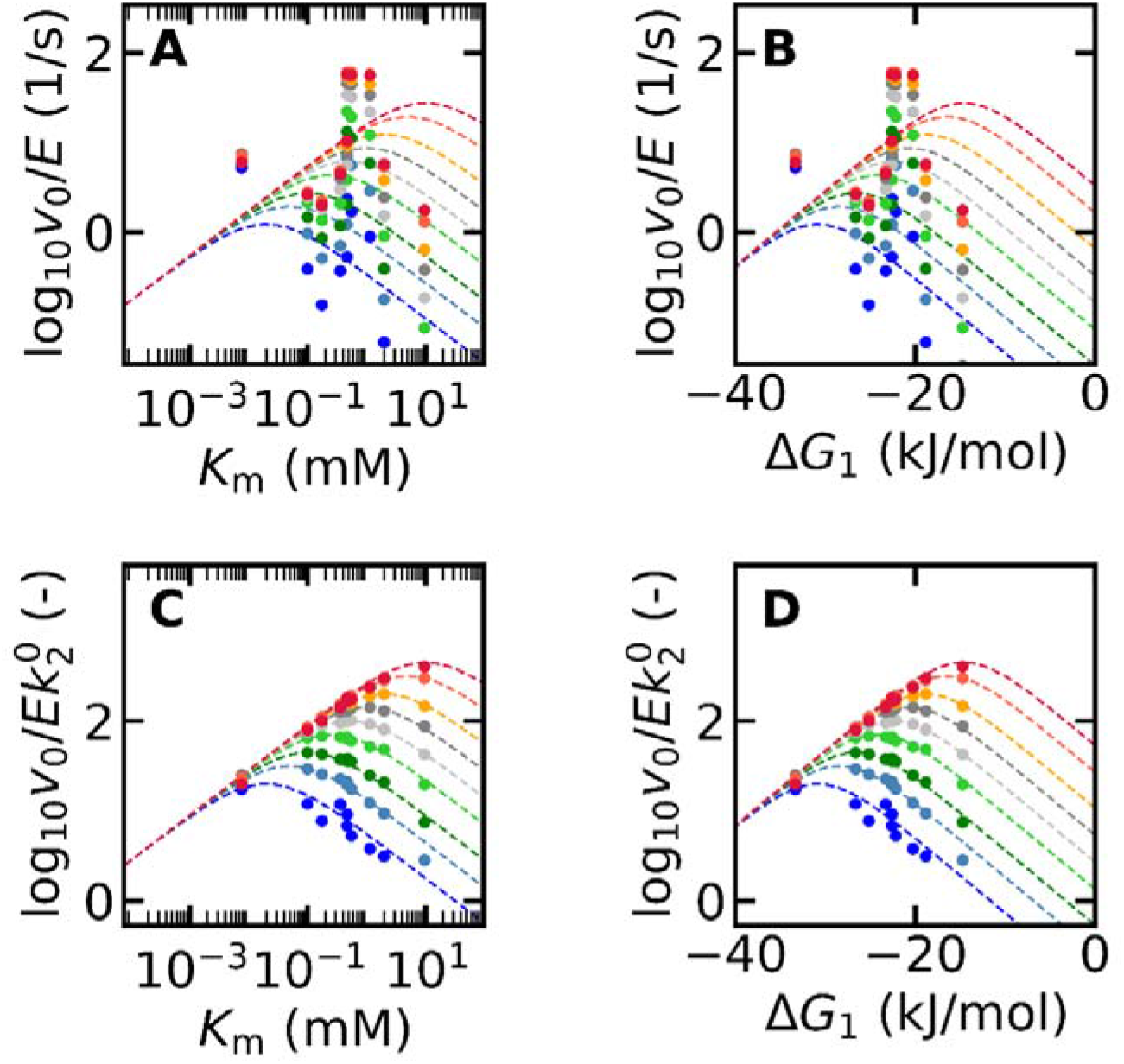
Relationship between the binding affinity (or) and the activity. (A) Markers indicate the raw data obtained from experiments and are the same with Fig. 2. Dashed lines indicate the theoretical activity calculated from the Michaelis-Menten equation (Eq. 3). (B) The X-axis of (A) has been converted to using Eq. 4. (C) The Y-axis of (A) has been normalized by calculated from Eq. 8. (D) Both X and Y axes have been converted using the raw data in (A).

## DISCUSSION

In this study we have demonstrated that the activity of PSP follows the Sabatier principle regardless of its phylogenetic and physiological diversity. PSP activity exhibited a local maximum at *K_m_* ≈ 0.5, which is conceptually consistent with the Sabatier principle in that activity is maximized at an intermediate binding affinity. Furthermore, PSP activity agrees quantitatively with the theoretical volcano plot after considering differences in 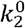, including the shift of the optimum binding affinity upon increasing the substrate concentration. Our findings indicate that both *K_m_* and 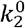 are necessary to rationalize PSP activity.

The influence of 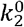 is not unique to PSP, because we found that normalizing the activity of cellulases by 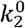 also results in a better match between experimental and theoretical activity (Fig S5). However, the influence of 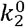 was not as pronounced in the case of cellulases due to its small variation. Namely, the 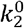 of PSPs were distributed across two orders of magnitude, while that of cellulase were within only one order of magnitude (Fig S6).

The reason for the different range of 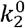 may be in part due to the diversity of our dataset; While the cellulases were obtained only from Fungi, our PSPs were obtained from all three domains of life. Furthermore, the natural operation temperature of each PSP is different, suggesting that each enzyme may have evolved under different physicochemical constraints. To compare the physicochemical parameters rigorously, we have intentionally determined the PSP activity at a constant condition (40°C, pH 8.0), irrespective of the natural operation environment of each enzyme. Their quantitative agreement with the activity calculated from the Sabatier principle under a condition completely independent from physiological context adds further support to the existence of an underlying physicochemical principle which dictates enzymatic activity.

While the conventional Sabatier principle attempts to rationalize enzymatic activity based on the binding affinity, expanding the theoretical framework by introducing 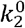 enhances its applicability towards a wider range of enzymes. Furthermore, explicit consideration of 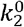 may also provide insight to realize enzymes with high activity. The correlation between *K_m_* and *k_cat_* reported by Kari et al (8) suggest that it is difficult to make an ideal enzyme with small *K_m_* and large *k_cat_*. However, if a large 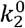 could be realized, *k_cat_* can be increased without increasing *K_m_*. Indeed, the *K_m_* and *k_cat_* of PSP does not show a linear relationship, due to the diversity of 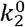 (Fig. S7). Although a direct strategy to manipulate 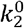 has not been found even for artificial catalysis, its diversity in PSP suggests that optimization may become possible in principle.

In conclusion, we have demonstrated that PSP activities can be rationalized based on the Sabatier principle, regardless of their sequences, three-dimensional structures, or phylogenetic and physiological background. Furthermore, PSP uses a soluble substrate, similar to the majority of existing enzymes, while cellulase and PET hydrolase have insoluble substrates. Just as the Sabatier principle has been expanded from heterogeneous to homogeneous catalysis, our results expand the scope of the Sabatier principle from heterogeneous to homogeneous enzymes, suggesting that an optimum binding affinity which maximizes the activity may exist for enzymes in general. Identification of an underlying physicochemical principle may be the key to understand the evolution of enzymes and metabolism, as well as design efficient enzymes in the future.

### Experimental procedures

#### Enzyme preparation

The PSP genes in *Meiothermus ruber* (K649_13560), *Chloroflexus aurantiacus* (Caur_0633), *Methanocaldococcus jannaschii* (MJ_1594), *H. sapiens* (PSPH), *E. coli* (SerB, JW4351), *B. subtilis* (BSU28940) were codon-optimized for *E. coli* and cloned between NdeI and EcoRI sites of pET21c. Codon-optimized PSP in *Persephonella marina* (PERMA_0343) was cloned into multicloning site 1 of pRSFDuet vector using NcoI and EcoRI sites. Previously synthesized expression vectors (the original gene sequences were used) were used for PSP in *H. thermophilus* (HTH_0103; multicloning site 1 of pCDFDuet) (20), *Thermosynechococcus elongatus* (TLR1532; pET21c) (21), and *Thermus thermophilus* (TT_C1695; pET26b) (23). The codon-optimized gene and amino acid sequences are listed in Supplementary Table S1.

*E. coli* BL21 Star (DE3) transformed with the expression plasmids were inoculated into Luria-Bertani medium containing the appropriate antibiotic(s): 50 μg ml ^−1^ ampicillin, 30 μg ml ^−1^ kanamycin, or 50 μg ml ^−1^ streptomycin. Cells were cultivated aerobically at 37 °C until the *A_600_* reached approximately 0.6, and then protein induction was performed by the addition of 0.5 mM isopropyl thio-β-d-galactopyranoside to the medium, followed by further cultivation for 3 h at 37 °C.

The harvested cells were resuspended in 20 mM Tris-HCl (pH 8.0) (buffer A), sonicated, and centrifuged to obtain the supernatant. The PSPs were purified from the supernatant and the purity was confirmed using sodium dodecyl sulfate-polyacrylamide gel electrophoresis (SDS-PAGE) and Coomassie Brilliant Blue staining as follows: The supernatant of the following proteins were heat-treated at 70°C (K649_13560, Caur_0633, TT_C1695, PERMA_0343) or at 80 °C (MJ_1594 and HTH_0103) for 15 min and centrifuged at 20,000 × g for 20 min, and the resulting supernatant were further purified using ResourceQ or MonoQ and Superdex 200 Increase columns. PSPH, JW4351, BSU28940, and TLR1532 were purified by using HiTrap Butyl HP, MonoQ, and Superdex 200 Increase columns without heat treatment.

#### Protein Assay

Protein concentrations were measured using a Bio-Rad protein assay dye (catalogue no. 500-0006). Bovine γ globulins was used as a standard.

#### Enzyme Assays

PSP activity was assayed by measuring the production of inorganic phosphate by using Malachite Green Phosphate Assay Kit (BioAssay Systems). The reaction mixture contained 100 mM Tris-HCl (pH 8.0 at room temperature), 1 mM MgCl_2_, 0.02-10 mM L-*O*-phosphoserine, and an enzyme solution with a total volume of 80 μl. The reaction mixture was routinely incubated at 40 °C or 70 °C for 10 or 15 min. The enzyme concentration and reaction time were adjusted so that the reaction speed was kept constant during the assay. The reaction was started by adding phosphoserine and stopped by placing the tube in ice-cold water, followed by the addition of highly acidic phosphate assay solutions. The inorganic phosphate concentration was determined by measuring the absorbance at 620 nm. The phosphate concentration in the reaction mixture without enzyme was subtracted as the background. One unit of activity was defined as the amount of enzyme producing 1 μmol of inorganic phosphate per min.

#### Mass Spectrometory

The protein bands separated by SDS-PAGE were cut out and destaind, followed by reduced alkylation with DTT and acrylamide. The samples were digested with trypsin (TPCK-treated, Worthington Biochemical) at 37°C for 12 hours. The resulting peptides were subjected to a MALDI-TOF MS (rapifleX MALDI Tissuetyper; Brucker Daltonics). The mass spectrometer was operated in the positive-ion mode and reflector mode the following high voltage conditions (Ion Source1:20.000 kV, PIE: 2.680 kV, Lens: 11.850 kV, Reflector 1: 20.830 kV, Reflector 2: 1.085 kV, Reflector 3: 8.700 kV).

The acquired data were processed using FlexAnalysis (version 4.0, Brucker Daltonics) and BioTools (3.2 SR5, Brucker Daltonics). The peptide mass fingerprinting was carried out with MASCOT (version 2.8.0, Matrix Science) against the in-house database including the amino acid sequences of HTH0103 and TLR1532 using the following parameters: enzyme = trypsin; maximum missed cleavages = 2; variable modifications = Acetyl (Protein N-term), Oxidation (M), Deamidated (NQ), Propionamide (C), Gln->pyro-Glu (N-term Q); product mass tolerance = ± 150 ppm.

### Mathematical calculations

Mathematical formulas were obtained by hand and Python 3.8.3 was used for the numerical simulations. *K_m_* and *k_cat_* were calculated using the *v*_0_ at substrate concentrations between 0.02 and 5 mM using a nonlinear model. A Δ*G_T_* = −50 kJ/mol was used based on the calculation using eQuilibrator 3.0 (27, 28) at 20 uM of phosphoserine, 1 uM of phosphate and 1 uM of serine at pH 8.0, pMg 3.0, Ionic strength 0.25M, and 25□.

### Derivation of Eq. 3

Below, we will denote *k_cat_* as *k*_2_ following convention in catalysis to clarify the similarity between *k*_1_ and *k*_2_. In order to estimate ΔG] from *K_m_* and *k*_2_, several assumptions were used. First, we have assumed that *k*_1*r*_ << *k*_2_, because the entire reaction is thermodynamically favorable (Δ*G_T_* ≈ —50 kJ/mol). Therefore,

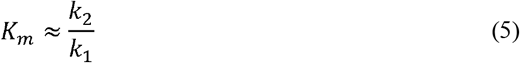

To bridge the driving force with rate constants, we have used the Arrhenius and Bronsted (Bell)-Evans-Polanyi (BEP) equations. For example, based on the Arrhenius equation, *k*_1_ can be expressed using the Arrhenius prefactor (*A*_1_), activation barrier (*E*_*a*1_), gas constant (*R*), and temperature (*T*) as:

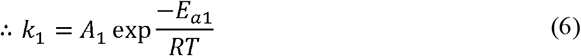

Based on the BEP relationship, activation barriers such as *E*_*a*1_ can be written as a function of the driving force as:

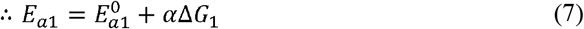

Here, *α* is the BEP coefficient and indicates how sensitive *E*_*a*1_ is with respect to Δ*G*_1_. In most cases, *α* > 0, because a more negative Δ*G*_1_, should promote the reaction by decreasing *E*_*a*1_. Simultaneously, it is common for *α* < 1, meaning that even if a driving force is applied, the activation barrier does not decrease by the same amount. Based on Eqs. 6,7, we obtain:

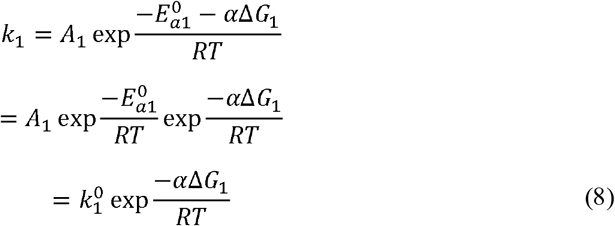

In the last row, we have defined 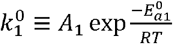 to highlight how *k*_1_ depends on Δ*G*_1_ Similarly, *k*_2_ can be expressed as

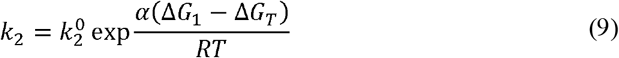

after taking into account Δ*G_T_* = Δ*G*, + Δ*G*_2_. Inserting Eqs. 8,9 into Eq. 5 gives:

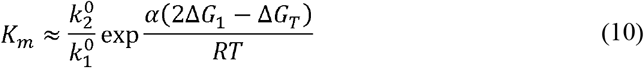

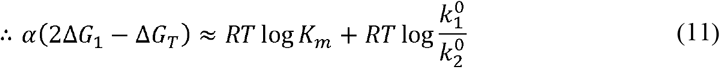

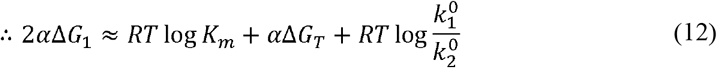

Assuming *α* ≈ 0.5, this yields Eq. 4 in the main text.

### Estimation of Experimental Parameters

Parameters such as Δ*G*_1_ and 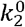 were estimated from the experimental data using a Python code which performs calculations using the derivations above. In order to determine Δ*G*_1_ using Eq. 4, several parameters must be obtained. *K_m_* was obtained directly from experiments, and Δ*G_T_* was set to —50 kJ/mol to reflect the experimental conditions in our enzymatic assay. Therefore, the bottleneck is to evaluate the ratio of the two equilibrium rate constants 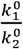. In this study, we have used the ratio of the Arrhenius coefficients 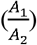 as an estimate. The Arrhenius coefficients were obtained as follows.

Based on Eq. 5, *k*_1_ was estimated for all enzymes as 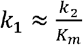. Out of the 10 enzymes in this study, 6 enzymes were thermostable at 70°C, and therefore, their Michaelis-Menten parameters (*K_m_* and *k*_2_) were obtained at 40 °C and 70°C. For these 6 enzymes, the Arrhenius equation was used to evaluate the prefactors (*A*_1_ and *A*_2_) along with their activation barriers (*E*_*α*1_ and *E*_*a*2_). The value of 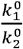 for all 10 enzymes was assumed to be the geometric mean of the value of 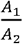 obtained from 6 enzymes. The accuracy of this approximation can be assessed based on the definition of 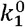 and 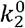. Namely, the BEP and Arrhenius coefficients are related as:

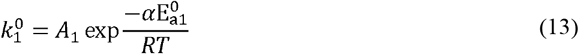

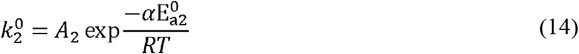

Therefore,

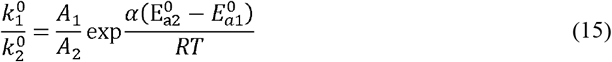

Thus, the approximation

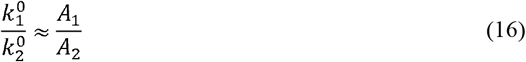

is valid when 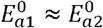. Although there is no experimental insight on the exact values of 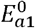 and 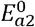, there is also no evidence which suggests one barrier is consistently larger than the other, and therefore, we have used Eq. 16 as a baseline assumption in our model. Deviations from this assumption will result in a mismatch between the experimental results and the theoretical line in Fig. 3. However, a 10 kJ/mol difference between 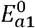 and 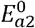 corresponds to a 7.4 fold deviation from Eq. 15. This is markedly smaller than the 2 order of magnitude variation within 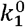 and 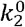, suggesting that the variation of 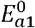 and 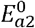 is not the main factor which dictates PSP activity.

## Data availability

Python code used to analyze the experimental data and export the figures will be made available at Github after publication (https://github.com/HideshiOoka/SI_for_Publications). The experimental assay data of PSP is available from the authors upon reasonable request.

The MALDI-TOF MS data used in the paper have been registered with PRIDE ARCHIVE (https://www.ebi.ac.uk/pride/archive) with the project accession number PXD040743 (Project DOI: 10.6019/PXD040743).

## Supporting information

Supporting information I.xlsx

Supporting information II.xlsx

## Acknowledgements

This work was supported by grants JPMJAX20BB (YC) and JPMJFR213E (HO) of Japan Science and Technology Agency (JST).

## Author contributions

Y. C., H. O., and R. N. designed the project. Y. C., M. W., and N. T., conducted enzymatic assays, N. D. and T. S. conducted mass-spectrometric analysis, and H. O. performed the mathematical calculations and numerical simulations. Y. C., H. O., and R. N. wrote the manuscript. All the authors have approved of the final version.

---

**Fig. S1.**
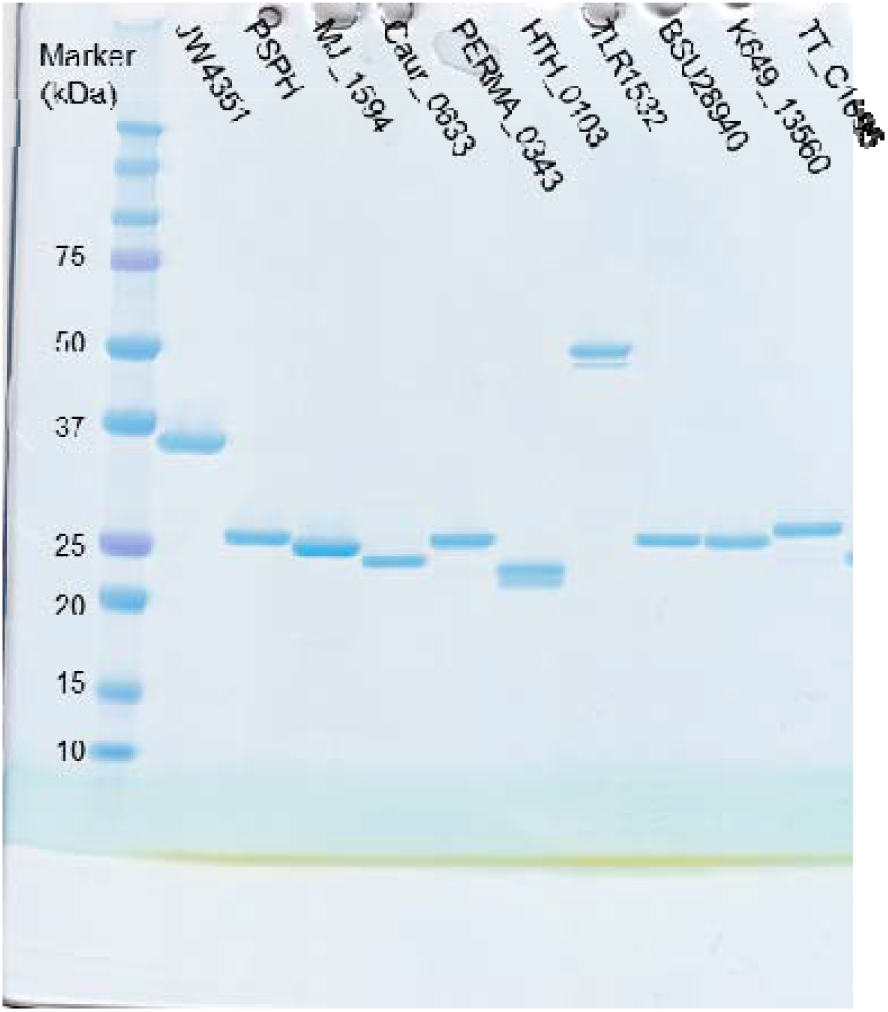
Purity of the enzymes used for the study. Approximately 2 μg of the enzymes were applied to each lane in SDS-PAGE and visualized by Coomassie Brilliant Blue staining. Mass spectrometry analysis confirmed that both major and minor bands observed in the lanes of HTH_0103 and TLR1532 are coding HTH_0103 and TLR1532, respectively, excluding the possibility of contamination of other proteins (see Supporting information II for details).

**Fig. S2.**
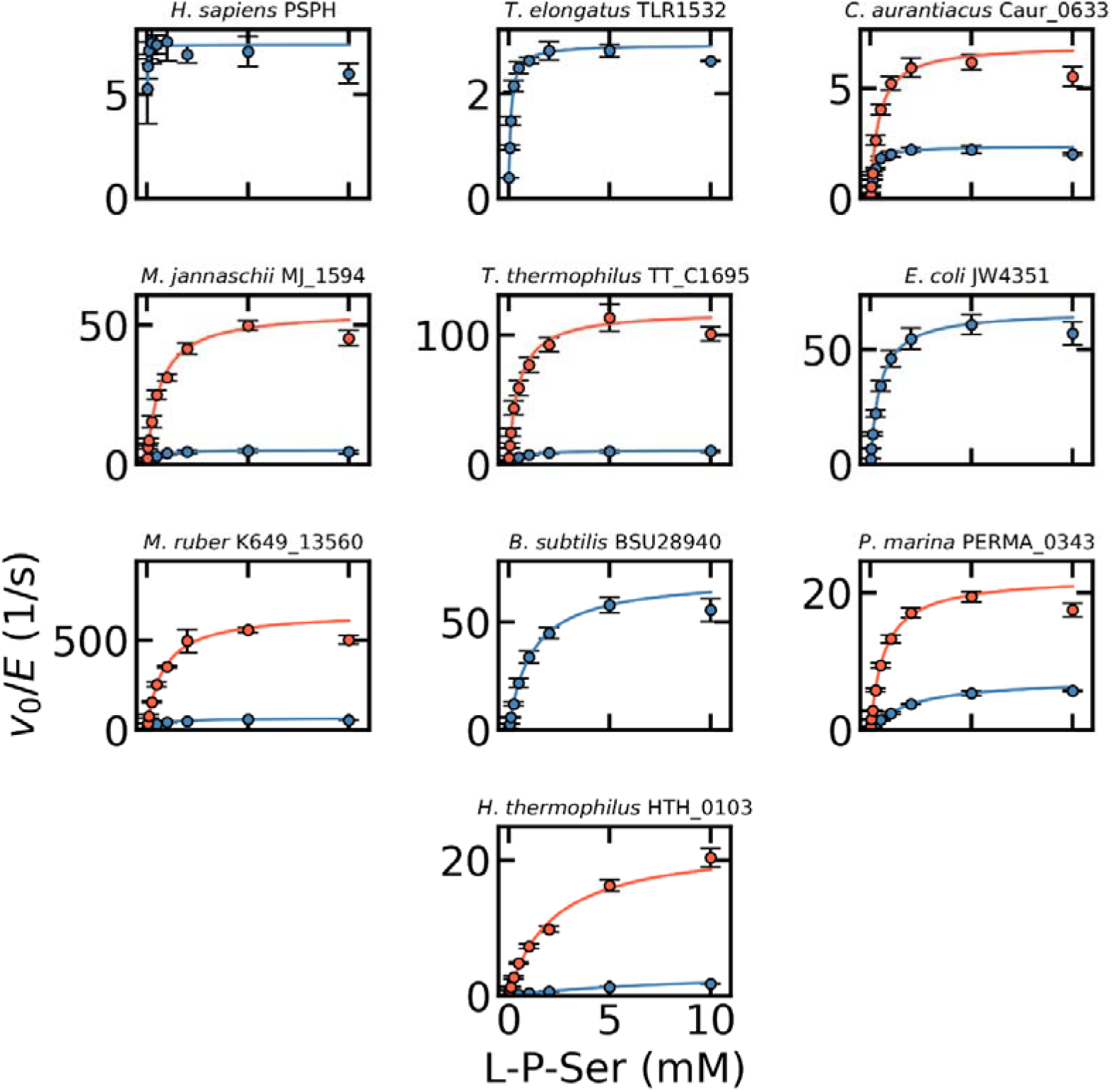
Michaelis-Menten plots of PSPs from variety of organisms. Orange and blue plots indicate the kinetics at 70□ and 40□, respectively. Error bars indicate their standard deviation.

**Fig. S3.**
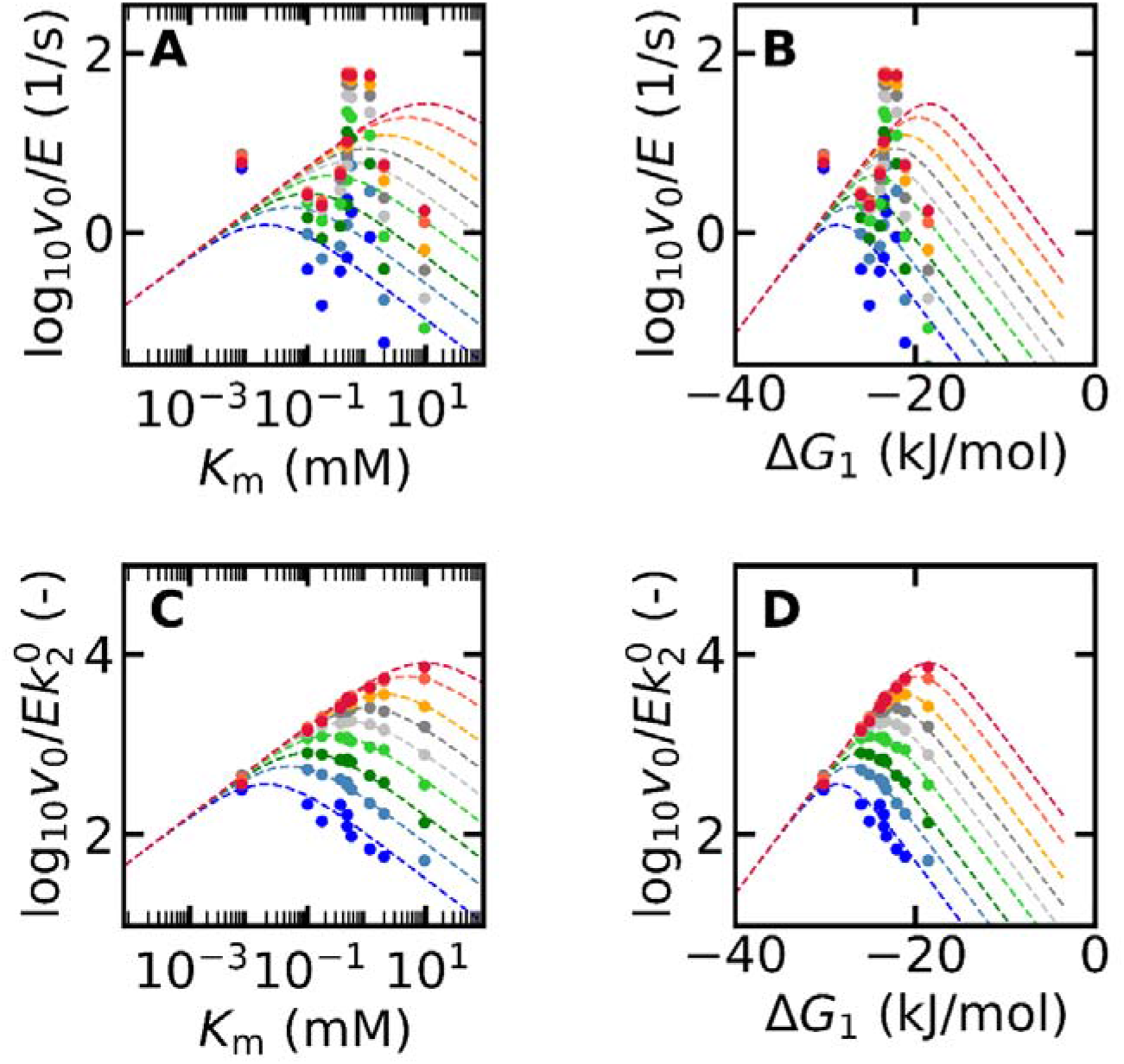
Influence of the BEP coefficient. The same analysis as Fig. 3 was performed using a BEP coeffiecient () of 0.8. This value was chosen based on the value reported for cellulases (0.74) [8]. The main conclusions, such as the observation of maximum PSP activity at a specific (A) and the quantitative agreement between theory and experiments upon normalizing by (C), are consistent regardless of the value chosen for the BEP coefficient.

**Fig. S4.**
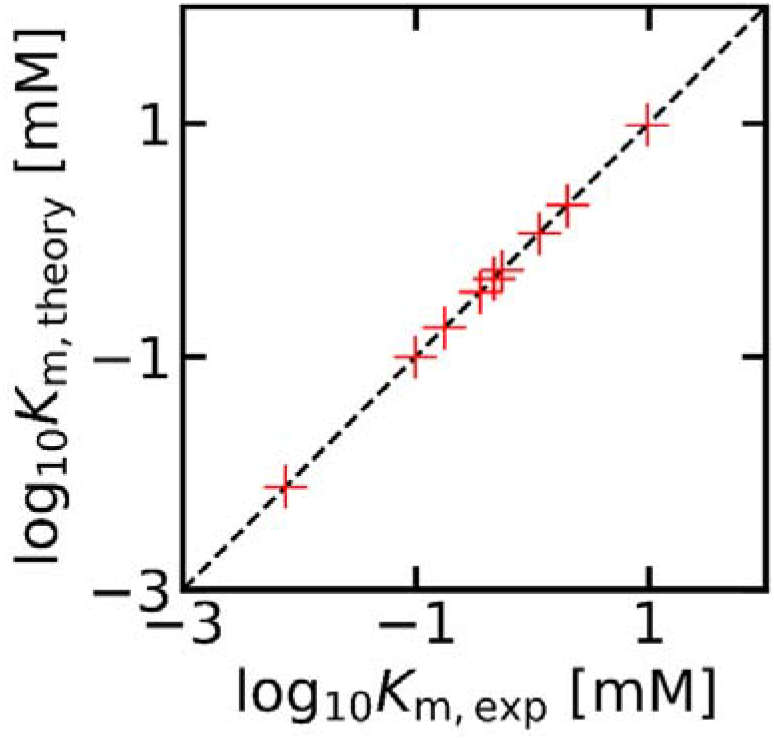
Comparison of experimental and theoretical values. The theoretical was calculated to confirm that the theoretically obtained parameters are reasonable. The value of was determined directly from experiments, and was approximated as (Eq. 4). Eq. 3 in the main text allows estimation of, and from this value, was calculated as —. The value thus obtained is approximately 4 orders of magnitude smaller than, and using the values of,, and thus obtained yield a theoretical value of which is consistent with the experimental value, showing that the approximation in Eq. 4 is self-consistent.

**Fig. S5.**
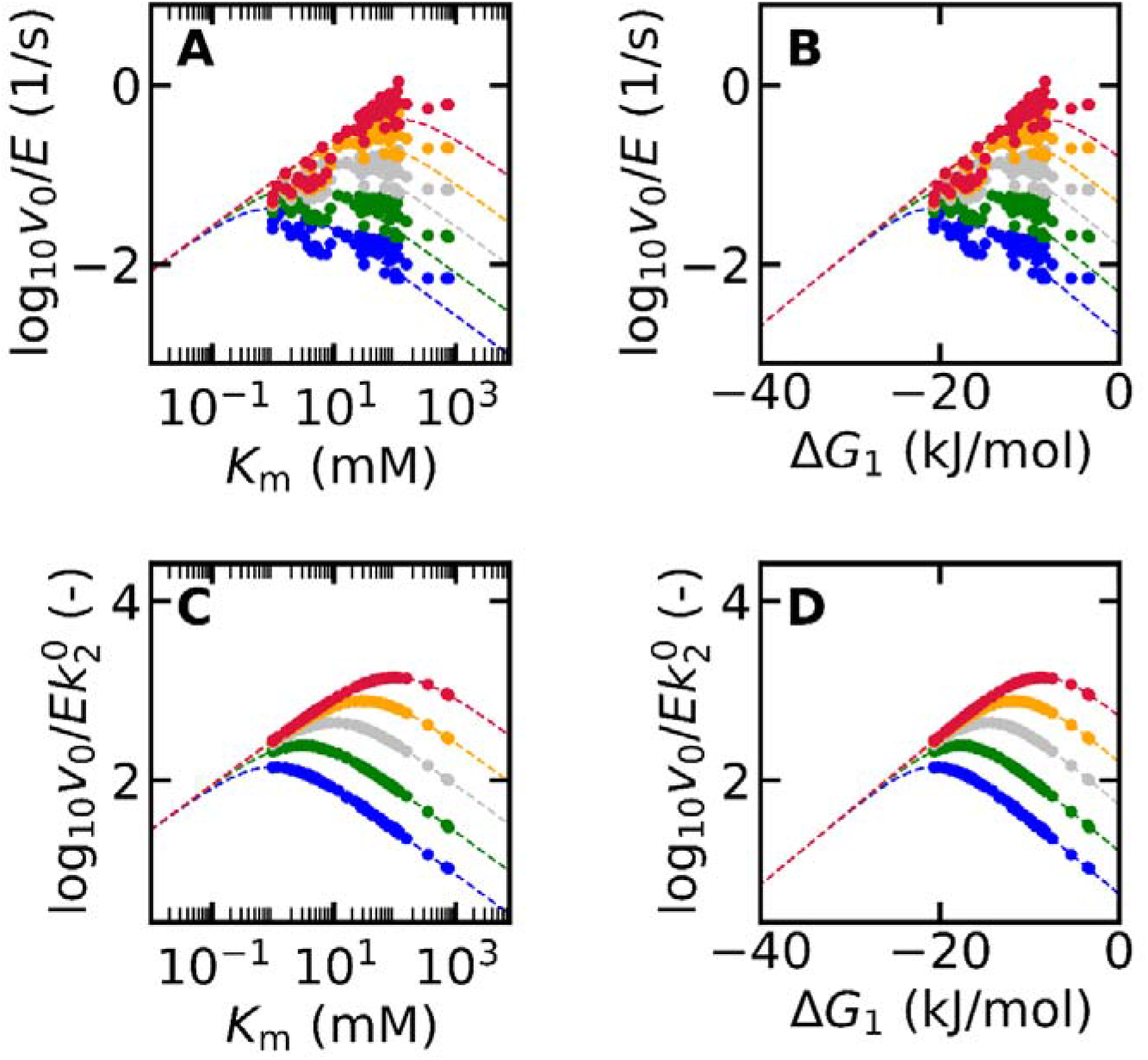
Volcano plots of cellulases. Raw data was obtained from ref. 8 and analyzed in the same way as Fig. 3 in the main text. Although the agreement between experiments and theory is reasonable without considering (A,B), normalizing with (C,D) markedly improves the agreement.

**Fig. S6.**
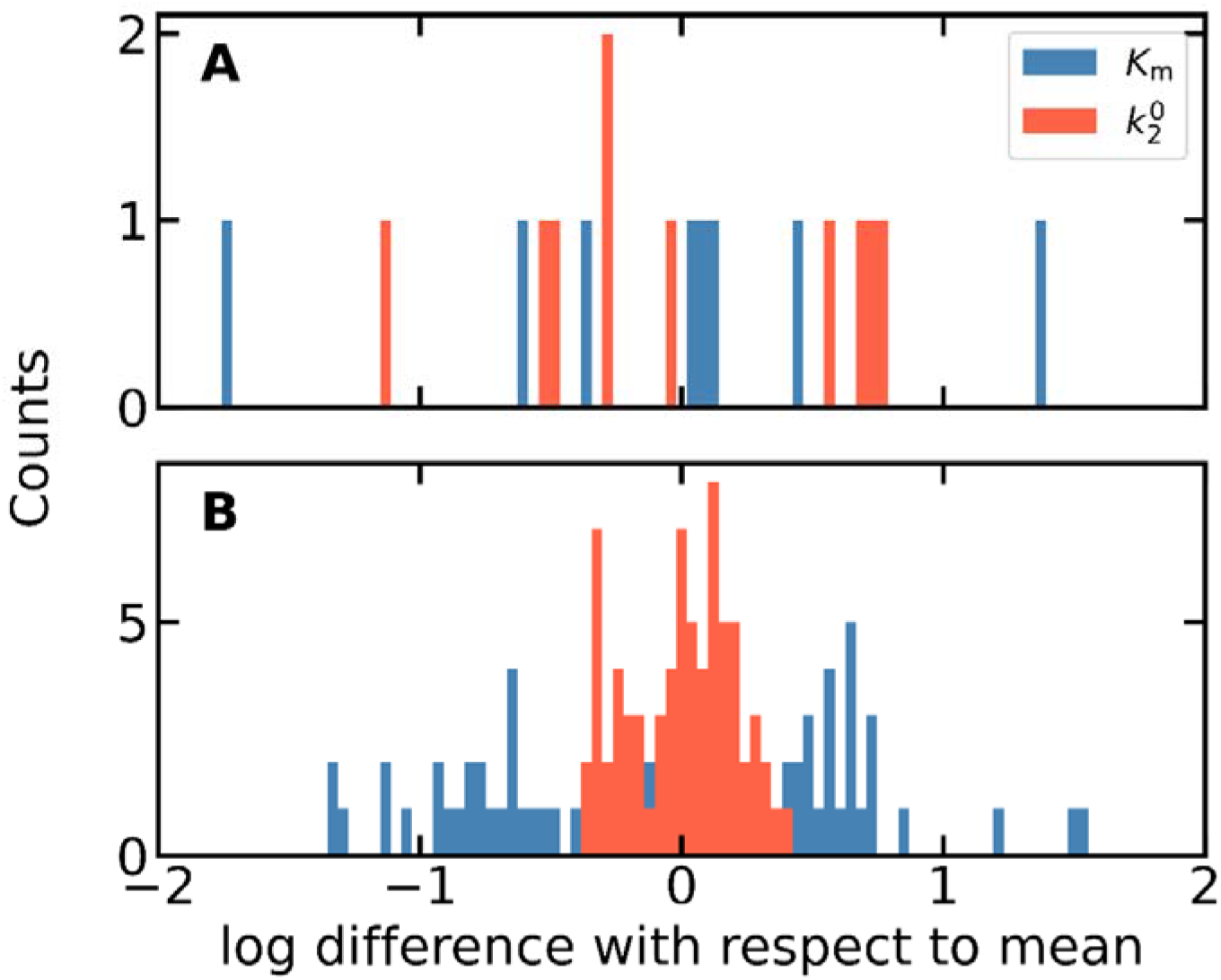
Diversity of and of PSP (A) and cellulase (B). The deviation of (blue) is of similar magnitude between PSP and cellulase. In contrast, the range of (orange) in cellulase is within an order of magnitude while that in PSP is almost three orders of magnitude. These results suggest that the influence of is larger for PSP.

**Fig. S7.**
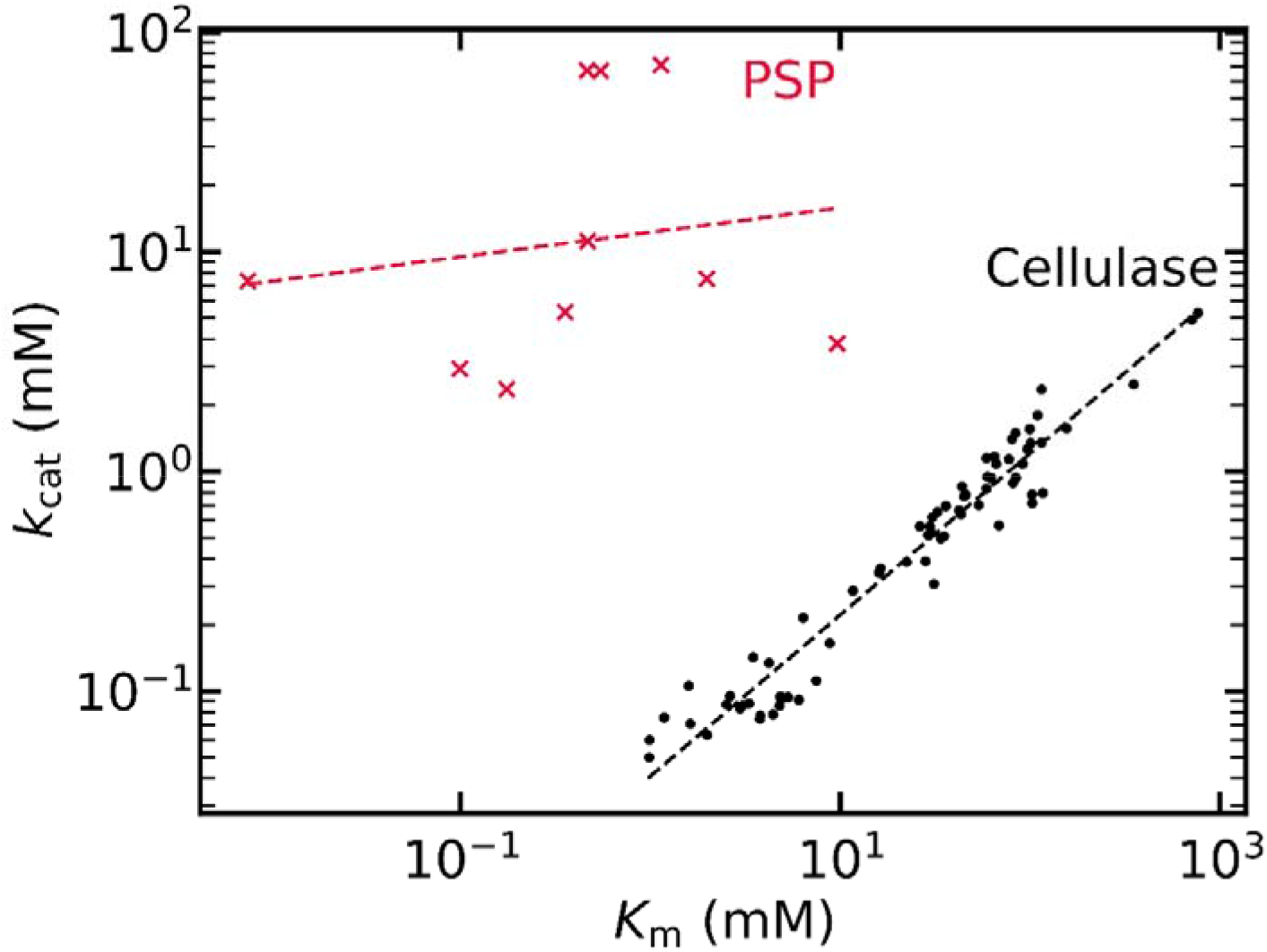
Scaling relationship between and. Cellulase data were obtained and replotted from ref. 8. The linearity in the case of cellulase is due to their small variation in. The mathematical relationship showing how influences the linearity between and is shown in Eq. 15 of ref. 22.

## References

1. Michaelis, L., and Menten, M. L. (1913) Die kinetik der invertinwirkung. Biochem. 49, 333–369

2. Johnson, K. A., and Goody, R. S. (2011) The original Michaelis constant: translation of the 1913 Michaelis-Menten paper. Biochem 50, 8264–8269

3. Michaelis, L., and Menten, M. L. (2013) The kinetics of invertin action FEBS letters,

4. Srinivasan, B. (2022) A guide to the Michaelis-Menten equation: steady state and beyond. FEBS J. 289, 6086–6098

5. Atkins, P., and De Paula, J. (2011) Physical chemistry for the life sciences, 2ndEd., Oxford University Press, USA

6. Kari, J., Olsen, J. P., Jensen, K., Badino, S. F., Krogh, K. B., Borch, K. et al.(2018) Sabatier principle for interfacial (heterogeneous) enzyme catalysis. ACS Catal 8, 11966–11972

7. Kari, J., Schaller, K., Molina, G. A., Borch, K., and Westh, P. (2022) The Sabatier principle as a tool for discovery and engineering of industrial enzymes. Curr Opin Biotechnol 78, 102843

8. Kari, J., Molina, G. A., Schaller, K. S., Schiano-di-Cola, C., Christensen, S. J., Badino, S. F. et al. (2021) Physical constraints and functional plasticity of cellulases. Nature Commun 12, 1–10

9. ArnlingBa□a□th, J., Jensen, K., Borch, K., Westh, P., and Kari, J. (2022) Sabatier Principle for Rationalizing Enzymatic Hydrolysis of a Synthetic Polyester. JACS Au 2, 1223–1231

10. Dyla, M., González Foutel, N. S., Otzen, D. E., and Kjaergaard, M. (2022) The optimal docking strength for reversibly tethered kinases. Proc Natl Acad Sci 119, e2203098119

11. Sabatier, P. (1911) Hydrogénations et déshydrogénations par catalyse Berichte der deutschen chemischen. Gesellschaft 44, 1984–2001

12. Ooka, H., Huang, J., and Exner, K. S. (2021) The sabatier principle in electrocatalysis: Basics, limitations, and extensions. Front Energy Res 9, 654460

13. Wodrich, M. D., Busch, M., and Corminboeuf, C. (2016) Accessing and predicting the kinetic profiles of homogeneous catalysts from volcano plots. Chem Sci 7, 5723–5735

14. Busch, M., Wodrich, M. D., and Corminboeuf, C. (2018) Improving the Thermodynamic Profiles of Prospective Suzuki-Miyaura Cross - Coupling Catalysts by Altering the Electrophilic Coupling Component. Chem Cat Chem 10, 1592–1597

15. Schramm, M. (1958) O-Phosphoserine phosphatase from Baker’s yeast. J Biol Chem 233, 1169–1171

16. Kuznetsova, E., Proudfoot, M., Gonzalez, C. F., Brown, G., Omelchenko, M. V., Borozan, J. et al. (2006) Genome-wide analysis of substrate specificities of the *Escherichia coli* haloacid dehalogenase-like phosphatase family. J Biol Chem 281, 36149–36161

17. Jung, T. Y., Kim, Y. S., Oh, B. H., and Woo, E. (2013) Identification of a novel ligand binding site in phosphoserine phosphatase from the hyperthermophilic archaeon *Thermococcus onnurineus*. Proteins 81, 819–829

18. Shetty, K. T. (1990) Phosphoserine phosphatase of human brain: partial purification, characterization, regional distribution, and effect of certain modulators including psychoactive drugs. Neurochem Res 15, 1203–1210

19. Jaeken, J., Detheux, M., Fryns, J. P., Collet, J. F., Alliet, P., and Van Schaftingen, E. (1997) Phosphoserine phosphatase deficiency in a patient with Williains syndrome. J Med Genet 34, 594–596

20. Chiba, Y., Oshima, K., Arai, H., Ishii, M., and Igarashi, Y. (2012) Discovery and analysis of cofactor-dependent phosphoglycerate mutase homologs as novel phosphoserine phosphatases in *Hydrogenobacter tthermophilus*. J Biol Chem 287, 11934–11941

21. Chiba, Y., Horita, S., Ohtsuka, J., Arai, H., Nagata, K., Igarashi, Y. et al. (2013) Structural units important for activity of a novel-type phosphoserine phosphatase from *Hydrogenobacter thermophilus* TK-6 revealed by crystal structure analysis. J Biol Chem 288, 11448–11458

22. Chiba, Y., Makiuchi, T., Jeelani, G., and Nozaki, T. (2016) Heterogeneity of the serine synthetic pathway in *Entamoeba* species. Mol Biochem Parasitol 207, 56–60

23. Chiba, Y., Yoshida, A., Shimamura, S., Kaineya, M., Tomita, T., Nishiyama, M. et al. (2019) Discovery and analysis of a novel type of the serine biosynthetic enzyme phosphoserine phosphatase in *Thermus thermophilus*. FEBS J 286, 726–736

24. [preprint] Ooka, H., Chiba, Y., Nakamura, R. (2023) Universal Design Principle to Enhance Enzymatic Activity using the Substrate Affinity. bioRxiv 10.1101/2023.02.01.526728

25. Ooka, H., and Nakamura, R. (2019) Shift of the optimum binding energy at higher rates of catalysis. J Phys Chem Lett 10, 6706–6713

26. Exner, K. S. (2020) Paradigm change in hydrogen electrocatalysis: The volcano’s apex is located at weak bonding of the reaction intermediate. Int J Hydrogen Energy 45, 27221–27229

27. Flamholz, A., Noor, E., Bar-Even, A., and Milo, R. (2012) eQuilibrator—the biochemical thermodynamics calculator. Nucleic Acids Res 40, D770–D775

28. Beber, M. E., Gollub, M. G., Mozaffari, D., Shebek, K. M., Flamholz, A. I., Milo, R. et al. (2022) eQuilibrator 3.0: a database solution for thermodynamic constant estimation. Nucleic Acids Res 50, D603–D609

